# Access to and use of preventive intermittent treatment for Malaria during pregnancy: a qualitative study in Chókwè district, Southern Mozambique

**DOI:** 10.1101/402727

**Authors:** Paulo Arnaldo, Isabel Cambe, Amílcar Magasso, Sérgio Chicumbe, Eduard Rovira-Vallbona, Anna Rosanas-Urgell, Sónia M. Enosse

## Abstract

**Background:** Malaria remains a significant health problem in Mozambique, particularly to pregnant women and children less than five years old. Intermittent preventive treatment is recommended for malaria prevention in pregnancy (IPTp). Despite the widespread use and cost-effectiveness of this intervention, the coverage remains low. In this study, we aimed to explore the factors limiting the access and use of IPTp-SP in Chókwè district.

**Methods and findings:** We used qualitative research methods through semi-structured interviews to collect data from 46 pregnant women and four health care staff from Chókwè, a rural area of southern Mozambique. Data were transcribed, manually coded and analysed using content and thematic method. Participants were not aware of pregnancy-related risks of malaria infection or the benefit of malaria prevention in pregnancy. Late and infrequently antenatal care (ANC) attendance, concerns about the long waiting time at ANC consultations,plus reluctance to disclose the pregnancy early, emerged as driving factors for inadequate IPTp delivery.

**Conclusions:** Pregnant women experience substantial barriers to receive adequate IPTp-SP dosing for malaria prevention. Poor awareness, non-compliance with ANC attendance and poor attitude of health care staff were main barriers to IPTp-SP delivery. There is a need to strengthen actions that improve awareness about malaria and prevention among pregnant women, as well as quality services across the ANC services in order to increase IPTp-SP uptake.

## Introduction

Pregnant women are at high risk of malaria and its adverse consequences. It is estimated that 5.5 million pregnancies occur annually in endemic areas, where malaria can be attributable of 10,000 maternal deaths [1,2]. Malaria infections during pregnancy are often asymptomatic, but can still cause adverse consequences for both the mother and her child, including maternal anemia, impaired fetal growth, premature and still-birth, low birth weight, or congenital malaria, which have been associated with a high risk for infant mortality and morbidity [1,3].

The control and prevention of malaria in pregnancy (MiP) is therefore an important strategy to avert malaria adverse consequences and improve maternal and infant health. The World Health Organization (WHO) recommends a package of intermittent preventive treatment in pregnancy (IPTp) with sulphadoxine-pyrimethamine (SP), use of insecticide-treated bed-nets (ITNs), together with effective case management of clinical malaria and anemia [3,4] in areas of high and moderate transmission in Africa. IPTp-SP together with ITNs is delivered during antenatal care (ANC) visits. Even though many countries in sub-Saharan Africa (SSA) have adopted IPTp-SP for malaria prevention in pregnancy, the coverage of the recommended three or more doses is still unacceptably low, despite a modest increase in ANC attendance among pregnant women [5], limiting the beneficial effect of this strategy on maternal and child outcomes [6,7].

Quantitative data collection approaches have been widely used to explore factors affecting access and use of IPTp-SP. In this regard, several factors such as limited access to ANC services, health professionals attitudes and practices, low awareness of malaria consequences during pregnancy, low patient adherence, or community attitudes towards preventive interventions have been associated with low IPTp-SP coverage [7–10]. However, quantitative data collection often lacks inclusion of sociocultural data, such as individual and community’s sociocultural beliefs, that may be affecting pregnant women’s access and use of malaria control intervention [10–13].

In Mozambique, policies to improve maternal and neonatal health, such as those targeting anemia and malnutrition, the prevention of MiP, increased institutional deliveries, delayed age of first pregnancy, and reduction of unsafe abortions have been implemented countrywide [14,15]. Since 2006, IPTp-SP is administered though the directly observed therapy approach, and ITNs are distributed free of charge to pregnant women at ANC visits [15]. However, according to the 2015 national survey data, the proportion of women receiving two doses and three or more doses of IPTp-SP was 34.2% and 22.4%, respectively [16], whereas our recent study in Chókwè rural district showed a coverage of 46.6% for three or more IPTp-SP doses at delivery [17].

This study explored perceptions, experiences, and behaviours of key informants (pregnant women and health staff) about the IPTp-SP intervention, in order to identify the individual and community’s sociocultural factors contributing to inadequate receipt of IPTp-SP during antenatal care visits. This information may help to improve the IPTp-SP uptake in the region and in the country.

## Materials and Methods

### Study site and population

The study was conducted in Chókwè district, Gaza Province, Mozambique. Chókwè district is situated in along the Limpopo River and its population belongs mainly to the Changana ethnic group. Their main economic activities are subsistence farming, large-scale rice production, livestock keeping, small business and migrant labour in South Africa. Around 135,616 habitants are under continuous follow-up through a health and demographic surveillance system (HDSS). The HDSS covers an area of approximately 600Km^2^within a 25Km radius of Chókwè City, and is run by the “Centro de InvestigaçÃo e Treino em Saúde de Chókwè” (CITSC), a clinical research center affiliated with the Instituto Nacional de Saúde - Ministry of Health. The HDSS routinely registers pregnancies, births, deaths, and migrations [Bonzela et al, in preparation]. There are two seasons, a hot and wet season (from November to April) and a dry and cool season (from May to October). Malaria transmission is perennial and occurs year-round, being more intense during the rainy season. *P. falciparum* is the predominant malaria parasite species [18].

At the time of data collection, the country had adopted the latest WHO policy recommendation f monthly administration of SP, recommending women to receive a minimum of three doses ofSP during the course of a pregnancy. The Health Centers at Chókwè District offers maternal and child health treatment and preventive services that include screening and treatment of syphilis, anemia, and urinary tract infections,anthelmintic, ferrous sulphate supplementation and folate tablets, vaccination for tetanus toxoid, and prevention of mother to child transmission (PMTCT) of HIV [15]. The prevalence of HIV in women of 15-49 years old was 28.2%in2015 [16].

### Study design and data collection

This was a descriptive qualitative study design used to explore the factors limiting the access to IPTp intervention for malaria prevention during pregnancy. Data were collected between March and April 2015 in the context of a study aiming to evaluate IPTp-SP uptake and pregnancy outcomes. Four primary health facilities, namely, Chókwè Health Center, Terceiro Bairro Health Center, Lionde Health Center and Conhane Health Center; located in the area under HDSS surveillance and primary providing healthcare were purposively selected for the study.

Within the selected primary health facilities, women who were pregnant and aged ≥15 years old were invited to participate in the study. To have additional information on the IPTp delivery from the provider side, health care staff were also included in the study. This was represented by one nurse at each health centre, who had been responsible for the ante-natal consultation for at least one year preceding the date of interview.

Interviews were conducted in a reserved room by an experienced male social scientist assisted by a female research officer specifically trained for this study. Interviewing sessions were conducted in Portuguese and/or in Changana (local language) depending on participants preference. Pregnant women responded a semi-structured questionnaire developed to explore (a) general perception about diseases affecting the population in the study area, (b) perceptions about malaria and IPTp-SP, and (c) experiences and perceptions of ANC service quality (Form S1). Health care staff were asked about (a) women attitudes towards IPTp use, and (b) challenges for IPTp-SP delivery (Form S2). All interviews were digitally recorded and lasted between 45 minutes and one hour.

### Transcription and data analysis

All components of the interview recordings that were in the local language (Changana) were translated to Portuguese language and transcribed *verbatim* and double-checked for accuracy before the analysis. Data were coded using pre-defined themes based on the original research questions. Manual analysis was used during thematic and content data analysis. The data analysis process involved: 1) familiarization with data through reading and re-reading of transcripts; 2) refining the themes by comparing codes with similar ideas. The headings used in the results and discussion section of this paper reflect the codes used for the analysis. Reporting of the study methods and results follow the consolidated criteria for reporting qualitative research [19].

### Ethics approval and consent to participate

The study was approved by the National Health Bioethics Committee (CNBS) (IRB 00002657). Administrative approval to conduct the study was obtained from the local health facilities and the Ministry of Health of Mozambique. With participants’ prior agreement, written informed consent was obtained and recorded prior to the interview. Women aged below 18 years, provided informed assent and their husbands, mothers or representatives provided informed consents. During transcription, names were replaced with codes to ensure anonymity and digital recordings were deleted once transcription and translation were completed and quality approved.

## Results

### Characteristics of study respondents

In this study, a total of 50 participants were interviewed: 46 pregnant women attending ANC services, and four health care staff (nurses) trained for maternal and child health care and preventive services, with secondary or higher education. Two out of the four interviewed health care staff had been working in the Maternal and Child Health Services for three or more years, while the other two had been working for one year. Pregnant women were aged between 15 and 44 years old (median 25.5 [22-32]). Most women (48.8%), had primary education level, 37.7% had secondary education level, and 15.5% had no formal education. Thirteen percent of the pregnant women were married, 37.7% single, and majority were in a marital union (51.1%).

#### *a.* General perceptions about diseases

When responding to the open-ended question about perceived illnesses affecting the population, pregnant women identified HIV/AIDS, malaria, tuberculosis and diarrhoea as the most frequent health problems in the area. Other diseases cited as prevalent in lower proportion among women of childbearing age were the “*Xibelekelo*” (native language) - described as a disease linked to the female reproductive system, specifically to the uterus causing intense pain and attacking women during the cold season-, and “*Xitsongua tsonguana*”, described as a disease affecting pregnant women characterized by convulsions and high blood pressure. According to local belief *“Xitsongua tsonguana”* is something that can pass from the mother to the baby, and in case this happens, the baby can be born with the “moon disease” described as a set of symptoms such as body pain, convulsions, fevers, constipation or cough, that the baby can manifest every time the full moon appears. Other diseases cited by participants were “*Dzedzedze*” (fever), stress, and diabetes.

#### *b.* Perceptions about malaria

The majority (80.4%, 37/46) of pregnant women perceived malaria as one of the most important diseases affecting populations, mainly in the hot and rainy season. High fever, chills, weakness, lack of appetite, body pain, tummy ache, sometimes vomits and joint pains were mentioned as the main symptoms. The most common treatment for malaria mentioned by the interviewed pregnant women was Quinine and Coartem.

> *“I know about muntzototo (malaria) is the main health problem during rainy season. I got muntzototo (malaria) more than one time, but every time, I received treatment”.* [Pregnant woman].

All interviewees were unanimous in characterizing malaria as a disease widely known which attacks everybody, and they perceived that mosquito bites can cause malaria. Other reported that cause of malaria were keeping dirty surroundings with standing water, that may increase mosquito population. The study participants were also aware of the consequences of malaria infection to human health (anemia and death), and perceived pregnant women and children as groups most at risk for malaria infection and adverse consequences. However, when asked about the specific deleterious effects of malaria during pregnancy, very few (10.9%, 5/46) mentioned the adverse effects of malaria to pregnant women, such as lose of the baby when the infected mother is left untreated.

> *“Pregnant women are the persons who can contract “muntzototo” (malaria) very easily, I do not know how to explain, but this is the period of conceiving another life”-* [Pregnant woman]

Regarding prevention, most pregnant women (89.1%) were aware that malaria is preventable and recognized that keeping the house cleaned and using mosquito nets could prevent it. In view of the study respondents, their beliefs about malaria prevention encompass hygiene and cleanliness of the environment and use of mosquito nets and does not include the use of medicines like tablets.

> *“We have to clean up where we live, especially where we sleep, windows must be opened in the morning and closed at night and people should sleep in the mosquito net”-* [Pregnant woman].

#### *c.* Perceptions of IPTp

More than half (56.5%, 26/46) of the interviewed women mentioned to have heard about drugs given at the ANC to prevent diseases in general during pregnancy, but most of them could not mention either the name of the drug nor could mention what the drug is for. They mentioned having received “comprimidos” (Portuguese term for “tablets”) to refer to the three drugs given to pregnant women during antenatal care consultations for malaria prevention, though very few could specify which tablet was. Very few women (10.8%) were able to name it as *Fansidar*^*®*^, a commercial name for Sulfadoxine-Pyrimethamine (SP). In order to further explore the topic of IPTp, the interviewer referred to SP as “the white tablets given to pregnant women and normally taken in front of the nurse”.

> *“I did not know what was for the pills I was given. They just give me three pills and said take it, and after they gave me “comprimidos vermelhos” (iron supplement), which serves to increase blood. I took what they gave me”-* [Pregnant woman].

#### *d.* Attitudes and challenges towards IPTp use during pregnancy

The attitude of pregnant women towards the access and use of IPTp were predominantly positive, as the majority of women (76.1%, 35/46) mentioned having taken at least once the malaria prevention tablets during antenatal care visits. However, pregnant women do not complete the recommended doses during the gestational period. This was reflected in their delayed care seeking practices, as observed by health care staff. Moreover, health care staff complained that although women do receive SP, most of them (71.7%, 33/46) receive the first SP dose very late at five to seven months of gestation, which does not allow enough time to complete the recommended dosage. Pregnant women usually attend to one single ANC visit and only receive the second dose at the time of delivery.

> *“Women come for the first ANC visit at thirteen weeks of pregnancy and may receive up to three doses, but in most of the cases, women books first visit with eight months. These will not complete the recommended doses and sometimes we have to give them the last dose at the maternity”-*[Maternal and child Nurse].

Health facilities staff also claimed that lack of educating materials such as pamphlets, pictures or figures on the consequences of MiP constitute a difficulty to inform future mothers on the beneficial effect of IPTp-SP and therefore, this affects the compliance.

> *“ It is difficult to notice that pregnant women who are at higher risk do not seek treatment or do not take the completed IPTp dosage. Perhaps if we had some illustrative pictures or images of a person who did not take complete treatment it would be helpful. I think that this it’s a bit of a perception because some do not complete the recommended dosage because they think the pills are strong, in this way even with our counselling, women will not complete the dosage”.-* [Maternal and child Nurse].

When those who have some information and having received IPTp were asked about the reasoning to have taken it, some of them revelled to have started taking IPTp-SP because they were sick and did not want the baby to be born sick, while others mentioned having taken the drugs to prevent the baby from diseases. However, they had no detailed knowledge about the recommended dosing scheme.

“*I went early to my first antenatal care visit to know if I had some diseases, make control and receive drugs for malaria prevention in order to protect my baby. We cannot stay for many months without preventing malaria because when you do it while you are already sick the child can suffer and the doctors cannot do anything”*- [pregnant woman].

Waiting time at antenatal consultations was mentioned to influence the demand of pregnant women for ANC services. This was particularly important for patients living in areas far from the health facilities, and women responsible for other children. A pregnant woman, who had the first antenatal visit between four and six months of gestation, said:

> *“ This [waiting time] makes it a bit difficult because I live far away from the health centre when I imagine that I will stay here for a long time I give up”-* [pregnant woman].
>
> *“The person [pregnant women] says that […] I cannot come to the hospital soon because it gets very crowded, so I prefer to come here just for delivery”-* [Maternal and child Nurse].

#### *e.* Experiences and perceptions of ANC service quality

The interviewees said to have had good interaction with health workers during ANC consultations. However, the majority of them mentioned that the information about malaria and other important health issues should be delivered in several different ways. This could be through counselling during the consultations inside the office and during the daily lectures at the start of working days held in the health centers.

> *“They said we have to come to the hospital for antenatal care and for counselling. I did the registration and tested for HIV and the result was negative. The nurse gave me all the information and gave me pills to take. She also gave me ferrous salt and told me to take it once a day, I also received a mosquito net”-* [pregnant woman].
>
> *“And those pills you just took them without asking what was for?”-* [interviewer]
>
> *“I only was given the pills and a took them. They did not explain anything to me, they just gave it and said take it”-* [pregnant woman].

When the health providers were questioned about the way in which the users were offered the anti-malarial drugs, they acknowledged that they could do counselling about malaria and its prevention during ANC consultations and the benefits of the drugs during pregnancy. However, they recognized that in most cases they only provide the pills without explaining the purpose and its benefits because the burden and number of patients seeking care is huge, and as a mechanism to manage such pressure, health providers are “forced” to perform a quick consultation.

> *“Usually, we are overworked, with much to do. And when the women come for a prenatal consultation, often I just give the tablets for malaria prevention, even without explaining carefully the details”-* [Maternal and child Nurse].

## Discussion

This qualitative study on the use of malaria prevention during pregnancy in Chókwè district identified five important themes: general perception of pregnant women about most common diseases in the study area, perceptions of malaria and IPTp-SP, attitudes towards IPTp-SP use during pregnancy and experiences and perceptions of ANC service quality. The semi-structured interviews with pregnant women and health workers provided useful information on the perceived gaps that challenges the uptake of IPTp-SP among pregnant women. The qualitative data supports quantitative findings from the study on IPTp-SP coverage among delivering women, that was undertaken in parallel to this study, reported similarly low levels of awareness and knowledge of IPTp services among pregnant women in the same area [17].

The results indicate that women recognized malaria as an important disease that affects the population and mainly pregnant women in the study area, particularly in the hot and rainy season. However, only a few appeared to understand that malaria in pregnancy can negatively affect the health of the fetus. This finding correlates with results from a previous study conducted in Manhiça district, southern Mozambique, in which pregnant women were not fully aware of malaria-associated adverse maternal and birth outcomes [10]. The observed low awareness about malaria risk and the adverse consequences of MiP, might negatively influence mothers adherence to IPTp-SP and general malaria preventive interventions. This calls for increased awareness-raising during ANC visits about the risk and consequences of malaria infection during pregnancy and the benefit of SP and other MiP prevention strategies [12,20]. However, a closer analysis of the contents and format of the health education sessions and other sources of information available in this community would be necessary to ascertain this link.

In this study, although pregnant women recognized the importance of preventing malaria, a minority appeared to fully recognize the use of medicines/tablets, such as IPTp-SP as a preventive mechanism for MiP. This is in agreement with previously reports from Kenyan pregnant women [21]. The results of the present study are considered robust, as women failed to mention IPTp-SP not only when discussing malaria prevention approaches in general, but also were probed about SP in particular. Similar findings were also reported in a systematic review in which the majority of women from southern African countries were found to be unaware of the use and benefits of IPTp-SP for malaria prevention during pregnancy [12]. It will be important to ensure not only that during ANC visits, women understand the importance of malaria prevention in pregnancy but also the role of IPTp-SP to reinforce the attitudes on IPTp-SP uptake. Furthermore, all opportunities to promote health in the field of malaria should address IPTp-SP.

Although most women mentioned having never heard about IPTp-SP, they remembered to have been given a drug during antenatal consultations. However, some could not remember the name of the drug given neither wether this was given to prevent malaria, but they were able to describe the three white tablets that were taken in front of the nurse. Previous studies in Africa identified perceived adverse reactions to the drugs as an important determinant factor for adherence [10,11,22]. In this study, there were no indications that perception of adverse reactions could have negatively influenced adherence since none of the respondents expressed concerns over the safety of IPTp-SP. On another hand, the long waiting time at ANC consultations and the absence of supporting materials such as illustrative pictures or images and pamphlets, that could be helpful to inform women during health education sessions, may influence the demand of pregnant women for attending ANC services and receive IPTp-SP as reported in other African settings [23].

Our findings on pregnant women attitudes towards IPTp-SP use showed that ANC attendance was likely to constitute barrier for an adequate IPTp-SP uptake. Nurses of maternal and health services interviewed in this study reported that pregnant women do not complete the recommended doses for the gestational period. This was mainly linked to late and infrequent ANC attendance, which led to missed opportunities for the provision of the recommended IPTp-SP dosage. This is in agreement with our previously reported results in the study on IPTp-SP coverage among delivering women conducted in the same setting [17]. Similar studies conducted in Uganda and Mali also found that late and infrequent ANC attendance were important factors influencing poor uptake of IPTp-SP and other malaria preventive measures [23,24]. Therefore, barriers to early ANC seeking behaviour are extremely relevant to the understand and improve the factors that influence IPTp-SP coverage. Some studies have reported a linkage between late and infrequent ANC attendance with pregnant women’s cultural beliefs. For example, reluctance to disclose their pregnancy early, the tendency to only starting to attend ANC when the tummy is visible, which in most of the cases happens after the fifth month and a trend to only seek help when sick [10,23,24].

Although the hospital is viewed as an entity of trust and obedience towards the health system in most of the communities [25], in this setting, the health care staff were viewed by the respondents as a “well-intentioned” during antenatal consultations. However, most of the pregnant women mentioned having simply taken the tablets without an explanation about what those tablets were for. This is consistent with previous reports in many settings in sub-Saharan Africa [21,26] in which poor attitude of health providers does influence adherence to ANC attendance and health interventions [11,27,28,29], and adds to the body of evidence calling for improvements in the quality of ANC services in the study area and in the country in general.

Our study findings should be interpreted taking into account the following limitations: the interview of pregnant women included in the study was performed among those attending ANC services and at the health facility. This may have limited the level of women comfort to openly give their opinions on the questions raised by the interviewer. In addition, conducting interviews in the community could have provided more insights from pregnant women not attending ANC consultations. A second limitation of the study rises on the inability of data collection through focal group discussion (FGD), which could have enabled us to explore experiences and beliefs concerning access and use of IPTp-SP among different population strata in the community. However, future research could explore further dimensions of factors influencing the delivery of preventive interventions for malaria in pregnancy.

## Conclusions

Our study findings highlight the importance of assessing the social factors that contribute to the access and use of IPTp-SP. The main factors influencing the access and use of IPTp for malaria prevention identified in this study were poor awareness about the risk and consequences of malaria infection during pregnancy, the added benefit of IPTp-SP for malaria prevention, non-compliance of antenatal care attendance, as well as the poor attitude of health care staff towards explaining IPTp-SP purpose. Actions needed to improve the implementation of MiP prevention strategies should include increased awareness about malaria and need of prevention among pregnant women through community health education sessions and other available sources of information and, importantly, improve the demand for quality services across the ANC services.

## Acknowledgements

The authors gratefully acknowledge all pregnant women and health workers who participated in this study as well as the ChÓkwè district authorities. We acknowledge the institutional support of the ChÓkwè Health Research and Training Centre (CITSC), the Instituto Nacional de SaÚde-INS and the Institute of Tropical Medicine Antwerp (ITM). We extend our profound gratitude to Dr Khátia Munguambe (Universidade Eduardo Mondlane, Faculdade de Medicina, Maputo, Mozambique) for her constructive comments and technical inputs. We also thank Celestino Sinai, who transcribed the study audio-recordings, Olga Manuel Ngoque and Pedro Baloi who contributed to the collection of data.

